# Direct Coupling Analysis of Epistasis in Allosteric Materials

**DOI:** 10.1101/519116

**Authors:** Barbara Bravi, Riccardo Ravasio, Carolina Brito, Matthieu Wyart

## Abstract

In allosteric proteins, the binding of a ligand modifies function at a distant active site. Such allosteric pathways can be used as target for drug design, generating considerable interest in inferring them from sequence alignment data. Currently, different methods lead to conflicting results, in particular on the existence of long-range evolutionary couplings between distant amino-acids mediating allostery. Here we propose a resolution of this conundrum, by studying epistasis and its inference in models where an allosteric material is evolved *in silico* to perform a mechanical task. We find four types of epistasis (Synergistic, Sign, Antagonistic, Saturation), which can be both short or long-range and have a simple mechanical interpretation. We perform a Direct Coupling Analysis (DCA) and find that DCA predicts well mutation costs but is a rather poor generative model. Strikingly, it can predict short-range epistasis but fails to capture long-range epistasis, in agreement with empirical findings. We propose that such failure is generic when function requires subparts to work in concert. We illustrate this idea with a simple model, which suggests that other methods may be better suited to capture long-range effects.

**Author summary:** Allostery in proteins is the property of highly specific responses to ligand binding at a distant site. To inform protocols of *de novo* drug design, it is fundamental to understand the impact of mutations on allosteric regulation and whether it can be predicted from evolutionary correlations. In this work we consider allosteric architectures artificially evolved to optimize the cooperativity of binding at allosteric and active site. We first characterize the emergent pattern of epistasis as well as the underlying mechanical phenomena, finding four types of epistasis (Synergistic, Sign, Antagonistic, Saturation), which can be both short or long-range. The numerical evolution of these allosteric architectures allows us to benchmark Direct Coupling Analysis, a method which relies on co-evolution in sequence data to infer direct evolutionary couplings, in connection to allostery. We show that Direct Coupling Analysis predicts quantitatively mutation costs but underestimates strong long-range epistasis. We provide an argument, based on a simplified model, illustrating the reasons for this discrepancy and we propose neural networks as more promising tool to measure epistasis.

## Introduction

Allosteric regulation in proteins allows for the control of functional activity by ligand binding at a distal allosteric site [1] and its detection could guide drug design [2, 3]. Yet, understanding the principles responsible for allostery remains a challenge. How random mutations dysregulate allosteric communication is a valuable information studied experimentally [4] and computationally [5]. Several analyses have highlighted the non-additivity of mutational effects or *epistasis*. This “interaction” between mutations can span long-range positional combinations [6], results in either beneficial or detrimental effects to fitness [7], and shapes protein evolutionary paths [8]. Given the combinatorial complexity of its characterization, empirical patterns of epistasis are still rather elusive [9–12]. Concomitantly, progress in sequencing has led to an unprecedented increase of availability of data arranged into Multiple Sequence Alignments (MSAs) [13] containing many realizations of the same protein in related species. Different methods have been developed to extract information from sequence variability, e.g. Statistical Coupling Analysis [14,15] was applied to allostery detection in proteins. It was argued that the allosteric pathway was encoded in spatially extended and connected *sectors*, groups of strongly co-evolving amino-acids, supporting that long-range information on the allosteric pathway is contained in the MSA. Another approach, Direct Couplings Analysis (DCA) [16], aims at inferring evolutionary couplings between amino-acids. Direct couplings predict successfully residue contacts [16] so to inform the discovery of new folds [17], allow one to describe evolutionary fitness landscapes [18, 19] and correlate with epistasis [20, 21]. In the context of allostery, there is no statistical evidence for the existence of long-range direct couplings that would reveal allosteric channels [22], in apparent contradiction with the existence of extended sectors reported in [15] and the observation of long-range epistasis [6].

In this work we propose a solution for this discrepancy, by benchmarking DCA in models of protein allostery where a material evolves *in silico* to achieve an “allosteric” task [23–29]. We consider recent models incorporating elasticity [24–27, 29], in which long-range co-evolution [26], elongated sectors [26] and long-range epistasis [29] are present and can be interpreted in terms of the propagation of an elastic signal [29]. We focus on materials evolved to optimize cooperative binding over large distances [27], and find four types of epistasis (Synergistic, Sign, Antagonistic, Saturation) that exist over a wide spatial range. We perform DCA and find that it predicts well mutation costs but is a rather poor generative model. Strikingly, it can predict short-range epistasis but fails to capture long-range one, in agreement with empirical findings [22]. We illustrate why it may be so via a simple model, which suggests that neural networks are better suited than DCA to capture long-range effects.

### Model for the evolution of allostery

We follow the scheme of [26, 27] where a protein is described by an elastic network of size *L* made of harmonic springs of unit stiffness (here we consider *L* = 12). Binding events are modeled as imposed displacements either at the “allosteric” or at the “active” site (each consisting of several nodes), as shown in color in Fig. 1A. Such imposed displacements elicit an elastic response in the entire protein and cost some elastic energy, which defines our binding energy (see Sec. 1 in S1 Text). Following [27], the fitness *𝓕* measures the cooperativity of binding between allosteric and active site and is defined as the energy difference *𝓕 ≡ E^𝒜c^ −* (*E^𝒜c,𝒜l^ − E^𝒜l^*) where *E^𝒜c^*, *E^𝒜l^* and *E^𝒜c,𝒜l^* are respectively the elastic energy of binding at the active site only (*𝒜c*), at the allosteric site only (*𝒜l*) and at both sites simultaneously (*𝒜c, 𝒜l*). The fitness can be rewritten approximately as (see Sec. 1 in S1 Text)

**Figure 1:**
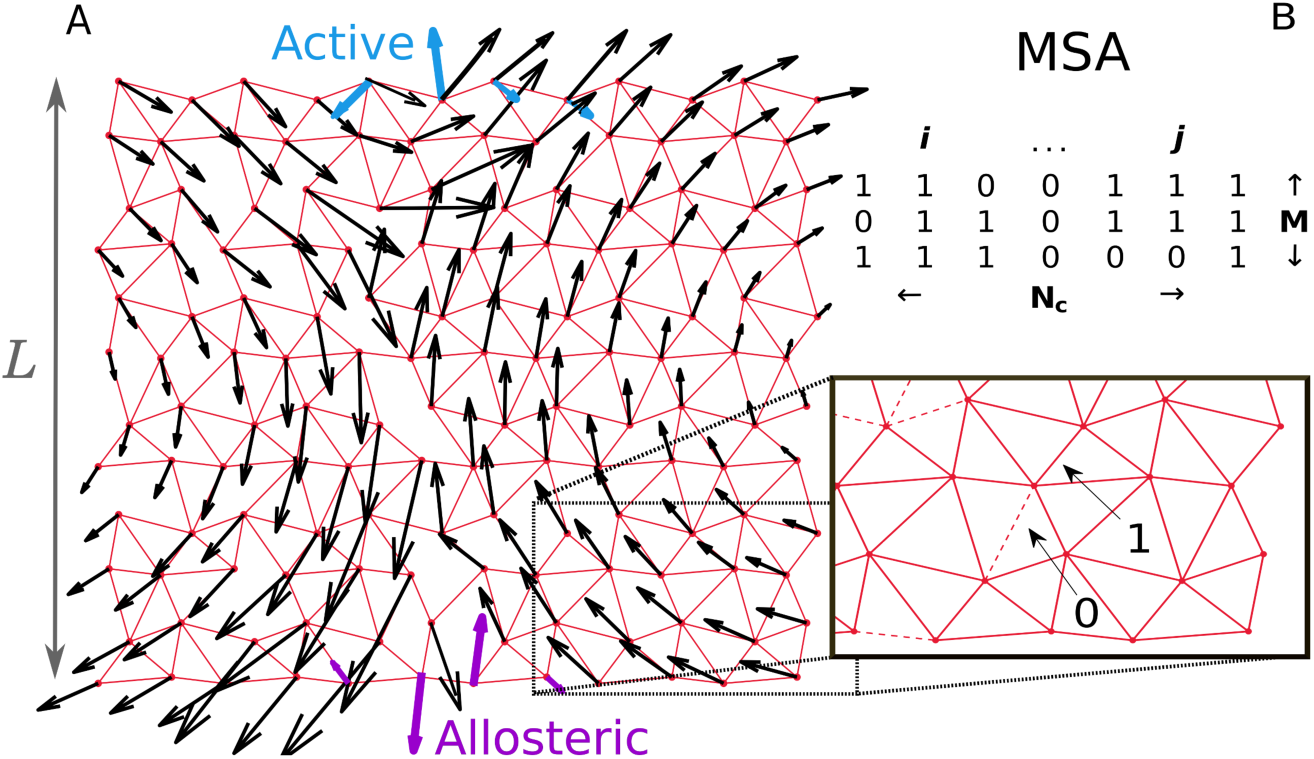
Study of co-evolution in artificial allosteric networks. A: Example of an elastic network made of harmonic springs (red) evolved *in silico* to maximize the cooperativity between the allosteric site (purple) and the active site (blue). The response to binding at the allosteric site is indicated by black arrows, and is found to follow a shear motion. B: Each network corresponds to a sequence of 0 and 1 coding for the spring absence or presence. Our scheme allows us to generate a large number *M* of such sequences, each corresponding to a slightly different shear architecture.

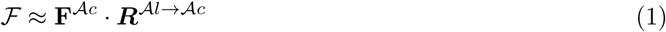

where ***F**^𝒜c^* is the force field imparted by substrate binding on the nodes of the active site, and ***R**^𝒜l→𝒜c^* is the displacement field induced at the active site by ligand binding. Note that each field in Eq. 1 is of dimension *n*_0_*d*, where *n*_0_ = 4 is the number of nodes in the active site and *d* = 2 the spatial dimension.

Such networks are evolved by changing the position of springs according to a Metropolis-Monte Carlo routine to maximize *𝓕*. At each step, the fitness difference with respect to the previous configuration ∆*𝓕* is computed and the new configuration is accepted with a probability *p* = min(1, exp *β*∆*𝓕*). *β* is an evolution inverse temperature controlling the selection pressure for high fitness *𝓕*, we choose *β* = 10^4^. We sample every 1000 time steps after an initial equilibration time of 10^5^ steps. At long times one obtains a cooperative system of typical *𝓕 ∼* 0.2, whose architecture depends on the spatial dimension and boundary conditions [27]. Here we consider a network in *d* = 2 dimensions with periodic boundaries, equivalent to a cylindrical geometry, where the response to binding evolves towards a *shear* mode. With our scheme we can generate thousands of networks with a similar design. A sequence ***σ*** of 0 and 1, where *σ_i_* = 1 stands for the presence of a spring at link *i* and *σ_i_* = 0 for its absence, can be associated to any network, leading to a Multiple Sequence Alignment (MSA) of networks performing the same function (see Fig. 1B).

## Results

### Nature and classification of epistasis

The cost of a single mutation (i.e. changing the occupancy) at some link *i* is defined as ∆*𝓕_i_* = *𝓕 − F_i_* where *𝓕* is the original fitness and *𝓕_i_* the one of the network after the mutation. We denote by ∆*𝓕_ij_* = *𝓕 − F_ij_* the cost of a double mutation at *i* and *j*. Epistasis between loci *i* and *j* is then defined as ∆∆*𝓕_ij_ ≡* ∆*𝓕_ij_ −* ∆*𝓕_i_ −* ∆*𝓕_j_*. Following Eq. 1 and observing that a mutation mostly affects the propagation of the signal ***R**^𝒜l→Ac^* and not how binding locally generates force (see Sec. 1 in S1 Text), epistasis follows approximately

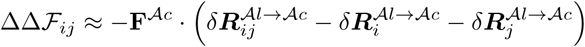

where 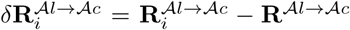, and 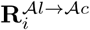 is the allosteric response at the active site of the protein mutated at link *i*. 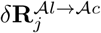 and 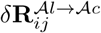 follow analogous definitions. We denote by *θ* the angle between 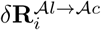 and 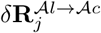. Assuming that the cost of a double mutation is dominated by the strongest point mutation, i.e. ∆*𝓕_ij_ ≈* max(∆*𝓕_i_,* ∆*𝓕_j_*) leads to

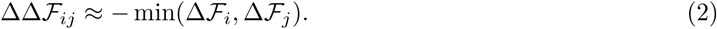

This assumption does capture a significant part of epistasis, especially when it is strong, as shown in Fig. 2A. This observation suggests to classify pairs of loci in terms of their epistasis and the minimal associated mutation cost min(∆*𝓕_i_,* ∆*𝓕_j_*) as performed in Fig. 2A.

*Saturation:* We define (somewhat arbitrarily) mutations with ∆*𝓕 >* 0.1 as lethal. Pairs of such lethal mutations (which represent *∼* 0.1% of all pairs, a sparsity in line with experimental findings [21]) have the strongest epistasis in absolute value, and follow closely Eq. 2, as visible in Fig. 2A. Physically, these mutations essentially shut down signal propagation by themselves with 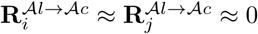, in such a way that the double mutation has the effect of a single one with 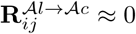. This view is confirmed in Fig. 2B by the observation that cos(*θ*) *≈* 1, as follows from 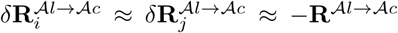. Saturation is then a form of very high “diminishing-returns” epistasis, for which evidence from data and support from theoretical models are accumulating [30, 31].

**Figure 2:**
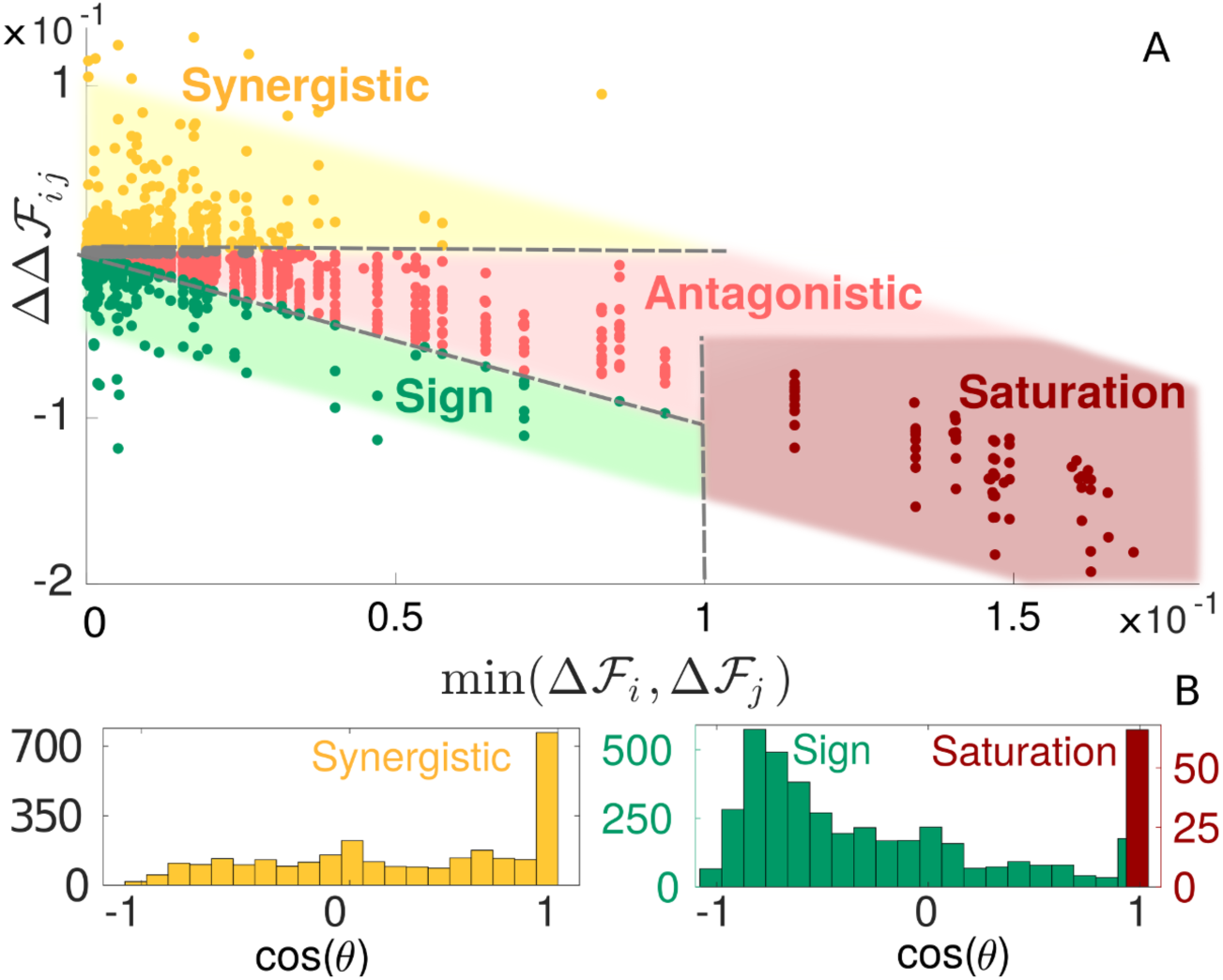
Classification and mechanical characterization of epistasis in our model of allosteric cooperativity. A: Phase diagram of epistasis in our allosteric material. All quantities are averages over 50 configurations obtained in a single run. The shaded area is taken with arbitrary width and a −1 slope as a guide to the eye. We show the lines ∆*𝓕_ij_* = 0, which divides synergistic from antagonistic/sign epistasis, ∆*𝓕_ij_* = max(∆*𝓕_i_,* ∆*𝓕_j_*), separating sign and antagonistic epistasis, and min(∆*𝓕_i_,* ∆*𝓕_j_*) = 0.1, the threshold set to distinguish lethal mutations. Points in grey correspond to epistasis *<* 5 *×* 10^*−*^4 and are excluded from our analysis. B: Histograms of *cos*(*θ*) for synergistic, sign and saturation epistasis.

*Antagonistic.* Further up along the diagonal of Eq. 2 in Fig. 2A, this saturation effect becomes milder. It is more akin to “antagonistic” epistasis [7, 32], whereby, after a first mutation, making a second one results only in a weak additional change.

*Sign.* In the intermediate range of negative-sign epistasis, more compensatory epistatic interactions can take place, where the fitness cost of a deleterious mutation is diminished by the second mutation (i.e. ∆*𝓕_ij_ <* max(∆*𝓕_i_,* ∆*𝓕_j_*)). Thus some mutations can become beneficial (i.e. increase the fitness) in presence of another mutation, and this resembles the “sign” epistasis empirically detected [7,33]. Geometrically, it corresponds to situations where the two mutations deform the signal in opposite directions, so the second one can partially re-establish fitness. In support of this, Fig. 2B shows that for sign epistasis cos(*θ*) tends to be negative.

*Synergistic.* Positive-sign values indicate “synergistic” epistasis. It occurs if two mutations perturb the elastic signal in the same direction, causing more damage than expected if they were purely additive. As clear from Fig. 2B, cos(*θ*) tends to be positive in this case.

### Direct Coupling Analysis

We evolve numerically *M* configurations maximizing cooperativity *𝓕*, each yielding a realization of a (variable) shear design. We sample a configuration for every initial condition to avoid introducing a bias in the sampling due to their high similarity. We find that the average Hamming distance among the obtained sequences is *∼* 20% of their length. Our set of sequences is analogous to a protein MSA – importantly, in this analogy the role of an amino-acid is played by a link, which can be stiff (*σ_i_* = 1) or not (*σ_i_* = 0, no springs). In practice we take *M* = 135000, much larger than the sequence length *N_c_* = (3*L*^2^ *−* 2*L*) = 408.

Next, for a statistical analysis of these sequences, we use DCA, which is based on the idea of fitting the observed single-site 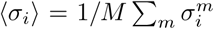 and pairwise 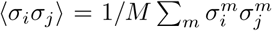 frequencies of links by the probability distribution *P* (***σ***) with maximal entropy (as this ensures the least biased fit of data under such empirical constraints). In our setup this approach leads to

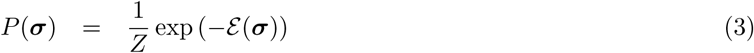

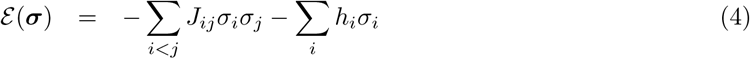

which is equivalent to an Ising model where *σ_i_* = 0, 1 would denote the two states (down, up) of spins. In this setting, *E* is an estimation of *βF*, *β* being the inverse evolution temperature. In all the comparisons (e.g. Fig. 3) we omit *β* as we are interested in the proportionality between *𝓔* and *𝓕*. The “fields” *h_i_* and “couplings” *J_ij_* are inferred to match *(σ_i_)* and *(σ_i_σ_j_)*. The inference of these parameters can be performed with several algorithms, we focus on ACE (Adaptive Cluster Expansion) [34, 35], an approximate technique developed from statistical physics ideas, combined with maximum likelihood, an exact technique. This approach is extremely accurate and we compare it to a method more approximate, but much faster computationally, as mean field Direct Coupling Analysis (mfDCA) [16], see Methods for details on the implementation.

**Figure 3:**
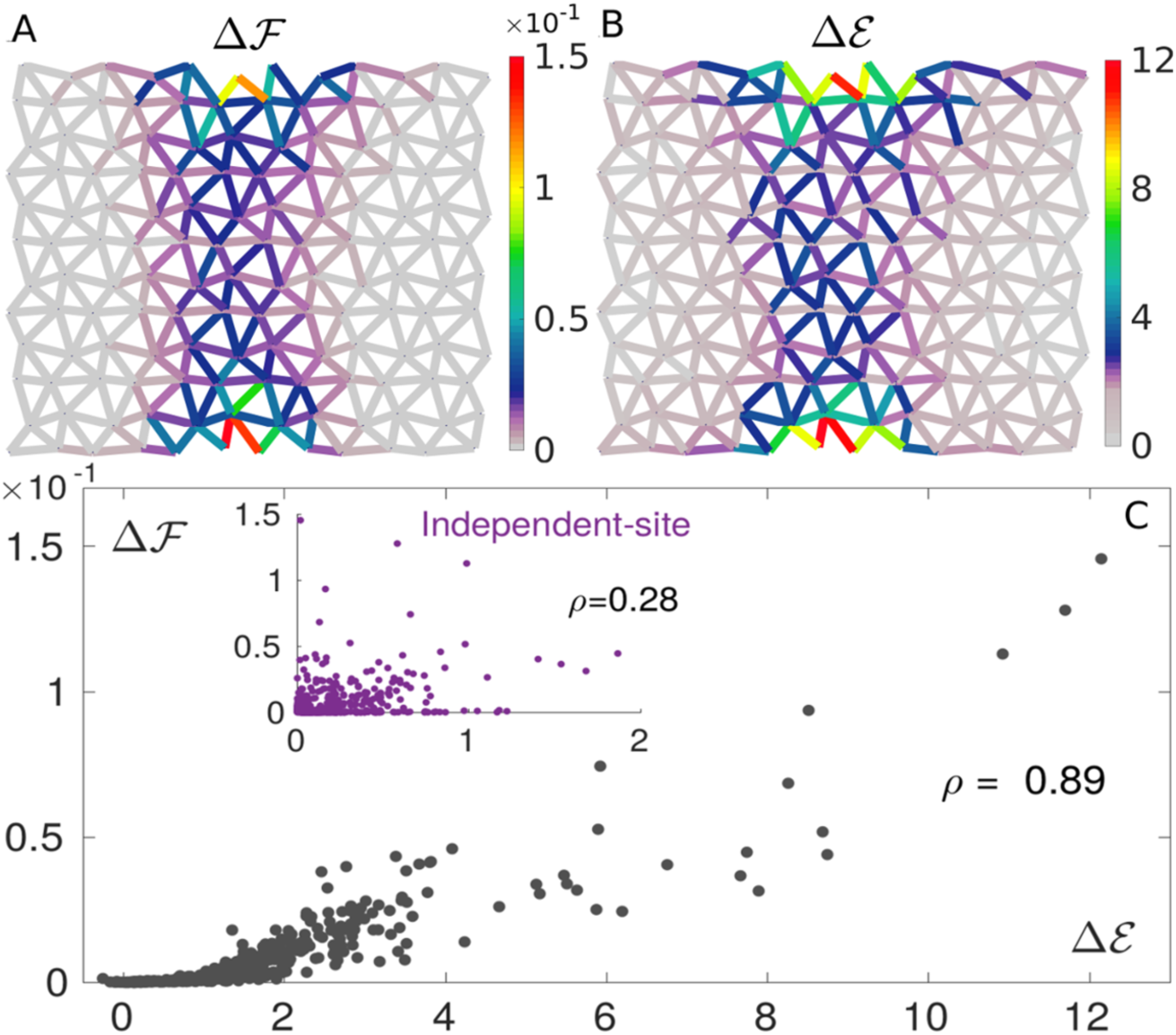
Prediction of mutation costs by DCA. Maps of true ∆*𝓕* (A) and DCA-inferred ∆*𝓔* (B) single mutation costs, averaged over 1.5 *×* 10^3^ configurations randomly chosen from the MSA. Their patterns are very similar, revealing high costs near the allosteric and active sites and in the shear path connecting them. C: Scatter plot showing the strong correlation between ∆*𝓕* and ∆*𝓔* for all links. The estimation of mutation costs based on an independent-site model (i.e. on conservation) correlates poorly with the true cost (inset), proving the need for incorporating correlations for proper prediction of mutation costs.

In this way we can benchmark DCA in the context of allosteric materials and test if it: (i) reproduces accurately the cost of single mutations; (ii) is a good generative model, i.e. if it can generate new sequences with high fitness and (iii) can predict epistasis.

### Inferring mutation costs

Fig. 3A shows the map of true mutation costs, indicating a large cost near the allosteric and active sites as well as in the central region where the allosteric response displays high shear (as documented in [27]). DCA enables one to infer this map by computing the estimated mutation cost ∆*𝓔_i_* = *𝓔_i_ − 𝓔* for a mutation at a generic link *i*, Fig. 3B. The comparison is excellent, as evident also from the high correlation revealed by the scatter plot Fig. 3C. Importantly, including pairwise couplings is key for inferring mutation costs, as a model based on conservation alone performs poorly, see inset of Fig. 3C.

### Generative power of DCA

Once the model of Eqs. 3, 4 is inferred, can it be used to generate new sequences with a high fitness, as previously shown for models of protein folding [36]? To answer this question, we generate new sequences by Monte Carlo sampling from the probability distribution Eq. 3. Fig. 4 shows the fitness of the obtained sequences vs their distance to “consensus” - the consensus being the most representative sequence of the MSA, i.e. where springs occupy the positions with largest mean occupancy. We find that (i) the variability of the MSA, quantified by the distance to consensus, is well reproduced (ii) the fitness is much more variable than for random sequences, with a few sequences that do perform as well as evolved ones (which never occurs for random sequences) but (iii) the mean obtained fitness is rather low, although larger, in a statistically significant way, than the one of random configurations (which is zero). As shown in Fig. 4, these results deteriorate further if a more approximate algorithm as mfDCA is used to infer parameters. We have checked that the generative performance is not improved by lowering the temperature of the

**Figure 4:**
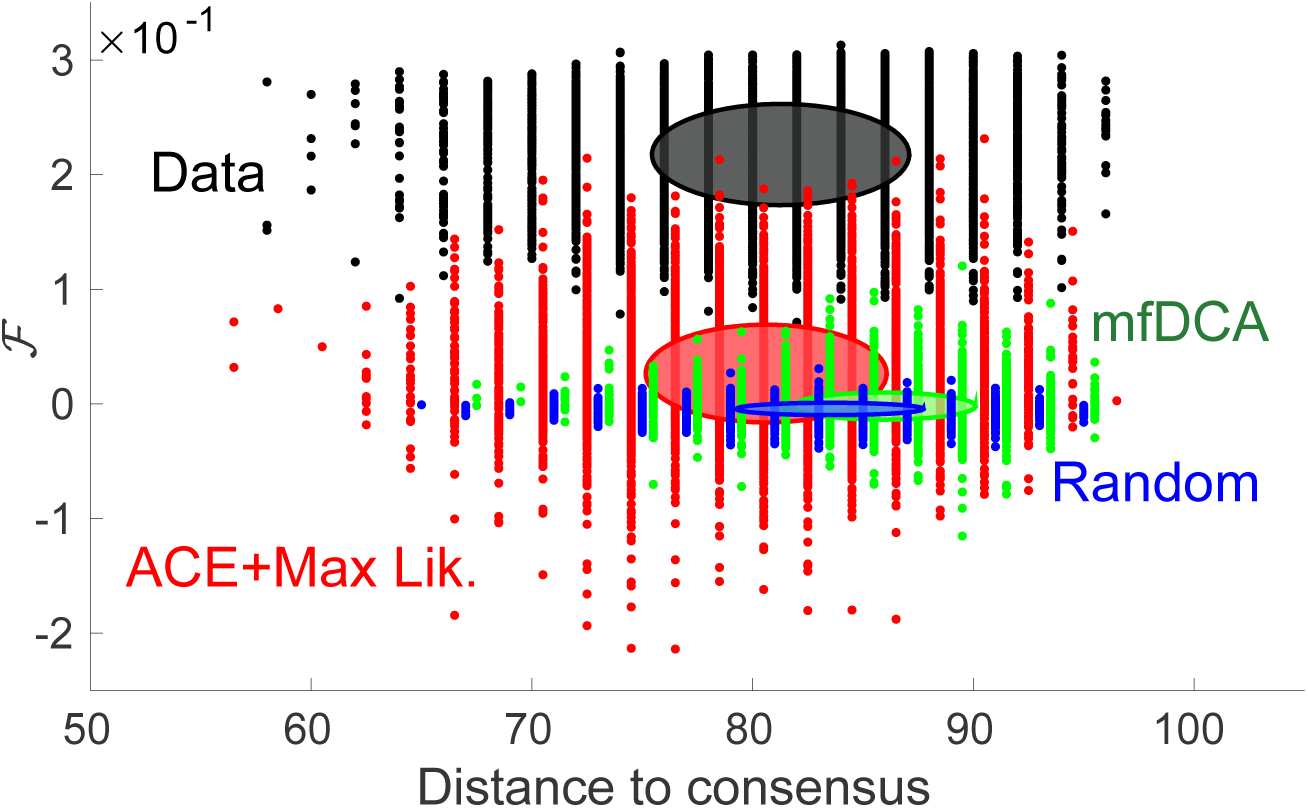
Generative performance of DCA. Fitness vs distance to consensus of configurations generated by the inferred model, following the representation of [36]. The sampling is done from *P* (***σ***) of Eq. 4 (a Boltzmann-Gibbs probability distribution), whose parameters have been inferred via ACE + maximum likelihood (red cloud) or mfDCA (green cloud). Original high fitness configurations (black cloud) and random ones (blue) are added as a reference. Each cloud consists of 104 sequences and the drawn ellipse gives one standard deviation around the mean in both horizontal and vertical directions. Distances to consensus of ACE + maximum likelihood, mfDCA and random sequences are shifted by respectively +0.7, *−*0.7 and *−*1.3 for better visibility.

Monte Carlo sampling. Overall, these results suggest that the generative power of DCA is limited in the context of allostery, in contrast with results for models of protein folding [36]. Thus an Ising model, a quadratic model accounting for conservation and correlations in the MSA (first and second order statistics), although it can capture some features of the shear design (e.g. the inhomogeneous distribution of coordination, as shown in Fig. S2), is a rather drastic approximation for the initial allosteric fitness. Indeed we have tested that higher orders as the third moment are not well reproduced (see Fig. S1). In what follows we shall emphasize in particular the failure of DCA to infer long-range epistasis.

### Inferring epistasis with DCA

From Eq. 4 one readily has that the DCA prediction for epistasis follows ∆∆*𝓔_ij_* = *−J_ij_*(2*σ_i_ −* 1)(2*σ_j_ −* 1), implying *|* ∆∆*𝓔_ij_|* = *| J_ij_|*. Hence, within DCA, the epistasis magnitude is simply the one of evolutionary couplings. In the inset of Fig. 5A we show the spatial location of the top 400 pairs of links with highest coupling magnitude, illustrating that long-range couplings are rare. Yet, as implied jointly by Fig. 2A and Fig. 3A, long range epistasis is present in our model, meaning that DCA fails to capture it. This fact is demonstrated quantitatively in Fig. 5A showing the mean epistasis *|* ∆∆*𝓕_ij_|* and mean DCA prediction *|* ∆∆*𝓔_ij_|* as a function of distances. The DCA-predicted trend reproduces the original one at small distances but strongly underestimates long-range epistasis. This is further evidenced in Fig. 5B showing that the average fraction of long-range pairs (range > 7) with the largest epistasis which falls in the list of the 400 pairs with largest couplings is much smaller than for short-distance pairs (< 7). However, even at short distance the prediction by *| J_ij_|* is not excellent, but it is remarkably improved if, as done in [12, 21], one considers epistasis averaged over several configurations (see Sec. 2 in S1 Text). Our finding is consistent with the lack of empirical evidence for long-range inferred couplings in allosteric proteins [22].

**Figure 5:**
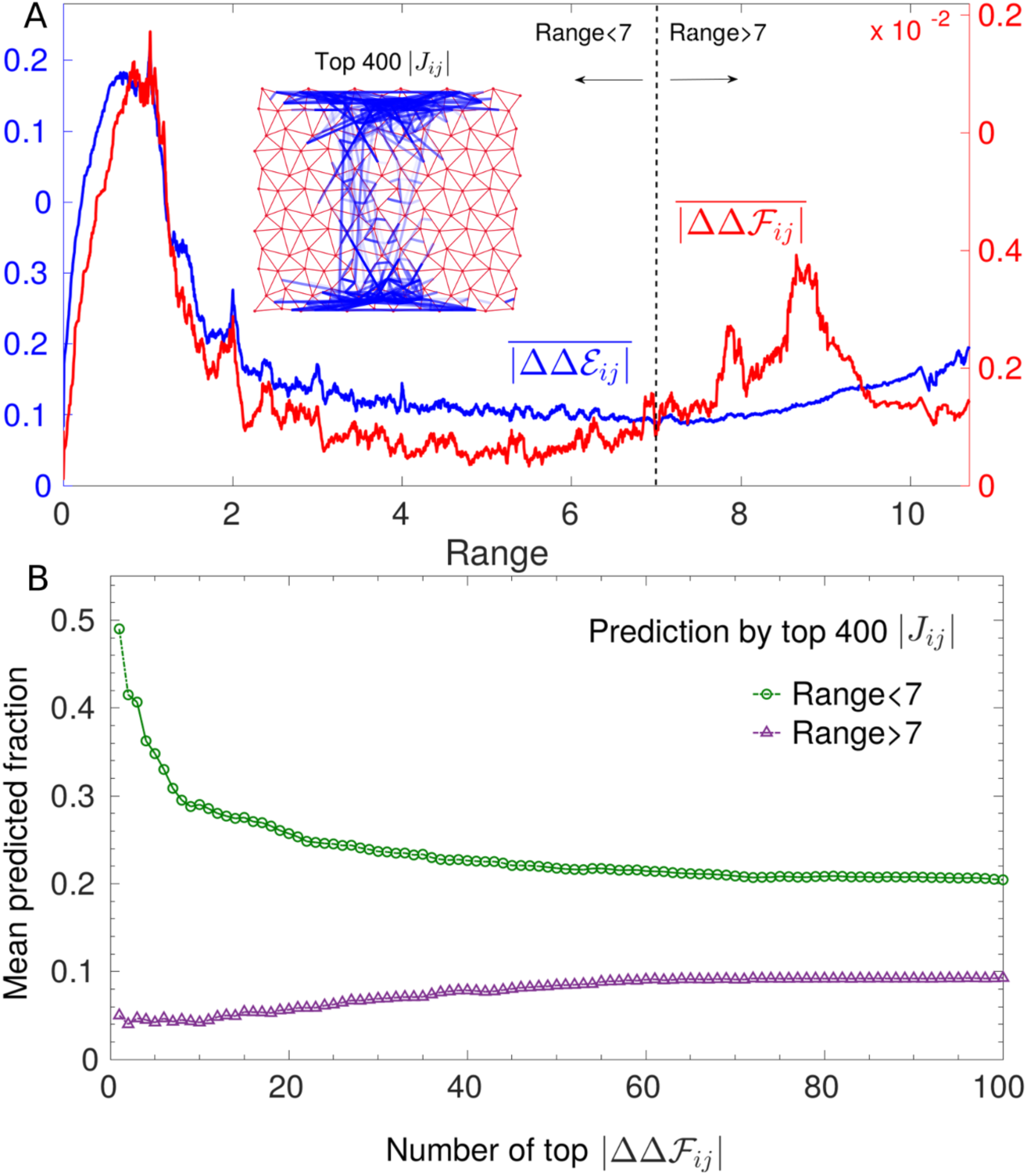
Prediction of epistasis by DCA. A: Running average of the absolute value of epistasis ∆∆*𝓕_ij_* and of DCA prediction ∆∆*𝓔_ij_* for 1.5 *×* 10^3^ configurations as a function of the distance between link *i* and *j*. The trends are nearly identical at short distances but at long distance DCA underestimates epistasis. Inset: Top 400 inferred couplings. They are mostly short range with only a few long-range couplings connecting the allosteric and the active site. Next we assess the prediction of epistasis in single configurations by these top 400 couplings. We consider separately long-range (> 7) and short-range (< 7) pairs of links, and rank them respectively in terms of the epistasis magnitude *|* ∆∆*𝓕_ij_|*. B shows which fraction of these pairs - averaged over 100 configurations randomly chosen - belongs to the 400 largest couplings, as a function of the number of pairs with maximal epistasis considered. Clearly coupling magnitude has less predictive power at large distances than at short ones. This feature stays robust also if we increase, e.g. up to 1000, the number of top couplings for prediction (see Fig. S4A).

### A proposed explanation for the failure of DCA at long-distances

We propose that the failure of DCA at long-range stems from its inability to describe a function that requires many subparts of the system to work in concert, when each subpart can be of different type. For example, in allosteric proteins on short length scales soft regions must exist where shear propagates [27, 37], giving rise to local constraints. Yet, there is flexibility in the exact location of these soft regions. On a larger length scale, these regions must assemble to create an extended soft elastic mode [27, 38, 39], which generates global constraints: for the shear architectures it implies the presence of a soft path between the allosteric and active site, whose position however can fluctuate. We argue that when applied to systems whose function is organized in such a hierarchical way, DCA underestimates long-range constraints. To illustrate this point, we introduce a Boolean model, shown in Fig. 6. A generic “function” is achieved by two subparts that must work in concert (AND gate) and that can be of two different types (OR gate) but each must be functional (AND gate). This model comprises 8 units, taking the value 0 or 1, decomposed into 4 groups: 2 groups are the possible types of subpart 1 (left in Fig. 6) and the other 2 the possible types of subpart 2 (right). A configuration is “functional” if 2 units of the same group are simultaneously in state 1 for each subpart. There are 49 functional configurations, whose fitness is fixed to *𝓕*, all other configurations have fitness 0. We assume that *𝓕* is large in such a way that the sequences in the MSA are only the 49 functional ones, with a uniform distribution. It is straightforward to calculate epistasis in this model, as well as single-site and pairwise frequencies from which couplings *J_ij_* and fields *h_i_* can be inferred. In particular we can compare ∆∆*𝓕_ij_* and ∆∆*𝓔_ij_* for units *i* and *j* either in the same group, so locally constrained by function (at “short distance”, e.g. *i* = 1 and *j* = 2), or in the two different subparts, thus globally constrained (at “long distance” e.g. *i* = 1 and *j* = 5). We obtain (see Sec. 2.1 in S1 Text) that *|* ∆∆*𝓕*_12_*| /|* ∆∆*𝓕*_15_*| ≈* 2.3: global and local constraints lead to relatively similar short range and long-range epistasis. Yet we find that epistasis between subparts is noticeably underestimated in contrast to epistasis within subparts. To show this, we look at the DCA prediction for the ratio of epistasis between two pairs of sites divided by the true ratio of epistasis. For pairs of sites belonging to the same subpart, DCA predicts equally well epistasis. For example, considering the pair of sites (1,2) and the pair (1,3), one finds *|* ∆∆*𝓔*_13_*| /|* ∆∆*𝓔*_12_*| × |* ∆∆*𝓕*_12_*| /|* ∆∆*𝓕*_13_*| ≈* 0.86 which is close to unity. However if sites belong to different subparts, DCA strongly underestimates epistasis with *|* ∆∆*𝓔*_15_*| /|* ∆∆*𝓔*_12_*| × |* ∆∆*𝓕*_12_*| /|* ∆∆*𝓕*_15_*| ≈* 0.33.

**Figure 6:**
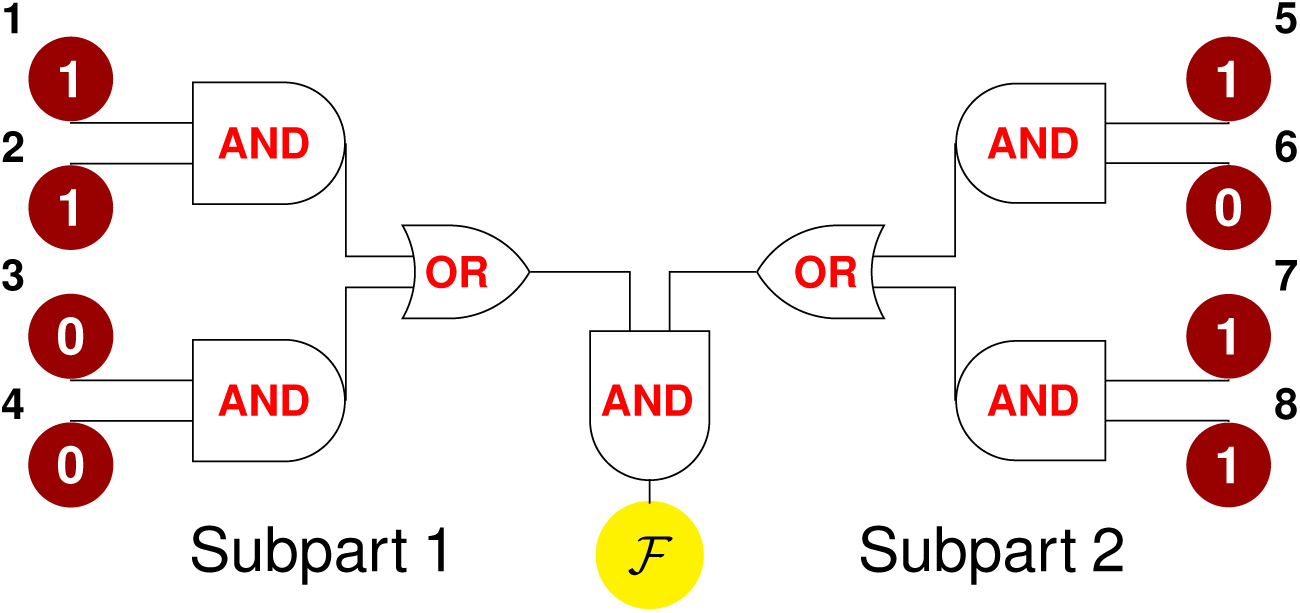
Sketch of a simple model for protein function. A system is arranged into 2 subparts which must work jointly to accomplish a given function (AND gate). Each subpart can be of 2 types (OR gate), to work each type must satisfy some constraints (AND gate between single units).

## Discussion

We have benchmarked DCA in a model of protein allostery where a mechanical task must be achieved over long distances. Such models display a rich pattern of epistasis, which can be both short and long-range and vary in sign. DCA predicts well mutation costs but is not a good generative model. This failure echoes with the drastic underestimation of long-range epistasis by the pairwise couplings inferred by DCA from evolutionary correlations. This finding rationalizes why there is no statistical evidence for long-range couplings in allosteric proteins analyzed by DCA [22], where long-range epistasis and functional effects are however found [6, 12, 15].

Yet, as we show in S1 Text (see Sec. 2), we expect that DCA can capture some aspects of the long-range epistasis pattern in allosteric proteins. Indeed, high-cost mutations exhibit stronger epistasis than low-cost ones (as also seen in RNA sequences [33, 40], in the enzyme TEM-1 *β*-lactamase [11] and in previous *in silico* evolution work [29]), and are well-predicted by DCA. Testing this DCA prediction for epistasis patterns empirically could be made possible by the increasing availability of deep mutational scans [12, 41].

Finally, we have provided the more general argument, illustrated by a simple model, that a co-evolution based maximum-entropy approach as DCA is not the appropriate inference framework when function requires several, variable parts to work in concert. Can one find better generative models than DCA for such complex functions? Several ways have been proposed to go beyond pairwise models by including nonlinearities, which implicitly take into account correlations at all orders, as nonlinear potentials in Restricted Boltzmann Machines [42], maximum-entropy probability measures with a nonlinear function of the energy [43] or maximum-likelihood inference procedures based on nonlinear functions [44]. As a first test, we have trained a 3-layers feedforward neural network with nonlinear (sigmoid) activation functions to learn the values of fitness in the simple model of Fig. 6. On the validation set, we could reach average mean squared errors on the estimated fitness *∼* 10^−6^ − 10^−7^, hence mutation costs and epistasis are correctly captured by this method (see Sec. 2.1.1 in S1 Text). This observation raises the possibility that neural networks may lead to better generative models in proteins, a hypothesis that could also be benchmarked *in silico*.

## Methods

### Direct Coupling Analysis: inference procedure

In a maximum-entropy approach, extracting information from MSAs can be cast as an inverse problem, i.e. inferring the set of parameters which enable the model (an Ising model in our setup) to reproduce certain observed statistical properties [45, 46]. The exact solution of this problem is found by Maximum Likelihood algorithms, which search for the set of couplings *J_ij_* and fields *h_i_* maximizing the likelihood that the model specified by such parameters produced data with the given statistics (single-site and pairwise frequencies in our case). This exact maximization might often be infeasible, therefore to tackle the inverse problem approximate techniques have been developed: for instance, we resort to the Adaptive Cluster Expansion (ACE), an expansion of the entropy (which indeed corresponds to the likelihood) into contributions from clusters of spins [34, 35, 47]. We use the package made available by Barton https://github.com/johnbarton/ACE. The implementation consists of first a run of ACE followed by a proper maximum likelihood refinement (QLS routine), which takes as starting set of fields and couplings the ACE-inferred ones. Different parameters for the ACE and QLS routines can be set by the user, e.g. *γ*_2_, the *L*_2_*−*norm regularization strength for couplings which penalizes spurious large absolute values induced by undersampling and for which a natural value is *γ*_2_ = 1*/M* (*M* being the size of the sample). To help convergence, we have chosen for ACE a higher value *γ*2 = 10^−2^ and *θ* = 10^−5^ (this is the threshold at which the algorithm will run then exit, see [35]). In the further refinement by QLS, we have set *mcb*, the number of Monte Carlo steps used to estimate the inference error, to 200000 and *γ*_2_ = 1*/M*. Having full control of the numerical evolution, we have tried to avoid undersampling issues by generating a large number of configurations *M* = 135000, which leads to *γ*_2_ *≈* 0.7 *×* 10^−5^. For the inference we remove from sequences the 6 links at the active and allosteric sites as they are always associated to the symbol 1 (always occupied by a spring), so the number of parameters to infer is 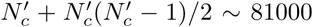 with 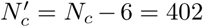. We have verified that low values of the *L*_2_-regularization allow us to obtain the maximal generative performance compatible with the model (in comparison to higher regularization). By default the *L*2 regularization of fields is 0.01 *× γ*_2_. In Fig. S1A, it is shown that the result of the inference is a model perfectly able to reproduce the first and second order statistics (as it should by construction) but that fails at reproducing higher order statistics.

For a comparison, we have considered also mean field Direct Coupling Analysis (mfDCA) [16], derived from a mean-field factorized ansatz for the Boltzmann-Gibbs distribution Eq. 3. Couplings in mfDCA are given by *J_ij_* = *−*(***C***^−1^)_*ij*_, where ***C**_ij_* = *(σ_i_σ_j_) − (σ_i_)(σ_j_)* is the covariance of the MSA (we recall that in each sequence *σ_i_* = 1 stands for the presence of a spring at link *i* and *σ_i_* = 0 for its absence). Typically ***C*** is not invertible due to undersampling, making it necessary to add a pseudocount *λ* (see [48]). As shown in [49], a pseudocount also helps correct for the systematic biases introduced by the mean field approximation: for this reason, we have used a pseudocount *λ* and chosen its value as *λ* = 0.5, which allows the best comparison to the ACE and maximum likelihood results, see Fig. S1B. It is noteworthy that in this way a computationally cheap technique as mfDCA yields a pattern of top *J_ij_* strikingly similar to the one of a very accurate inference achieved by the combination of ACE and maximum likelihood. Therefore mfDCA, while extremely poor as a generative model, exhibits a good performance at reconstructing the distribution of relevant couplings, as shown in Fig. S1C.

### Mutation costs and generative performance in the inferred Ising model

Costs of double mutations, i.e. joint mutations affecting links *i* and *j*, can be computed in the original model via fitness changes ∆*𝓕_ij_* = *𝓕 − 𝓕_ij_*, where *𝓕_ij_* is the fitness after springs in *i* and *j* have been mutated. A double mutation can correspond either to (i) adding two springs at links *i* and *j* (i.e. *σ_i_* = *σ_j_* = 1) or removing them (i.e. *σ_i_* = *σ_j_* = 0) or to (ii) moving a spring from link *i* to link *j* or viceversa (i.e. *σ_i_* = 0, *σ_j_* = 1 or *σ_i_* = 1, *σ_j_* = 0). Let us call the former “non-swap” mutations and the latter “swap” mutations. Swap mutations conserve the total amount of springs (360), thus the overall average coordination *⟨z⟩* = 5, and are the ones performed in the *in silico* evolution. As optimal allosteric configurations maximize fitness with respect to this type of mutations, we stick to them also when we compare mutation costs in terms of fitness and inferred energy (see Fig. 3C): we define “effective” single mutation costs ∆*𝓕_i_* and ∆*𝓔_i_* by taking, for each link, the swap with a link in the external region (more rigid, as visible in e.g. Fig. S2), where mutations are completely neutral, thus whose cost would be roughly zero.

For the generative step, we implement a Monte Carlo sampling which relocates springs from an occupied to an unoccupied link, i.e. which follows swap-type dynamics as for the original numerical evolution. This allows us to select, from the inferred model, sequences that are structurally as close as possible to the initial data, i.e. with the same average coordination *⟨z⟩* = 5, to make a consistent comparison with them. We have verified that even relaxing this constraint in the sampling leads to sequences endowed with higher internal variability yet lying in the same range on fitness (hence the inferred model incorporates rather well the information on the fixed amount of springs). The parameters of the Ising model are inferred in such a way as to match single-site occupancy, which reflects the spatial pattern of coordination in the allosteric networks. In Fig. S2 we show that generated sequences, despite having lower fitness, reproduce successfully this property as they should.

### Comparison with conservation

Single-site frequency in protein alignments, informative about local conservation, is a standard measure of mutation costs at a certain position [50] and can be fit by an independent-site Ising model. Energy (Eq. 4) in this case contains only field terms and, once these are inferred from link occupancies *⟨σ_i_⟩*, one can compute energy changes ∆*𝓔_i_* upon point mutations. The energy cost of a mutation in an independentsite model is then ∆*𝓔*_*i*_ = (2σ_*i*_ − 1)*h_i_*, where 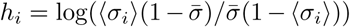 describes how the observed occupancy of a link *i*, *⟨σ_i_⟩*, is biased away from the average occupancy 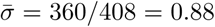. In average ∆*𝓔_i_* gives also a measure of *conservation* of link *i* as it is 0 when 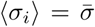 and it increases the more link *i* tends to be either occupied or vacant. The improvement achieved by the pairwise model over this conservation-based measure of mutation costs is extremely significant (see inset of Fig. 3C). On the one hand, conservation is a purely local measure - it takes into account how a particular position is crucial to the propagation of the allosteric response. Including pairwise couplings proves to be crucial to capture the context-dependence of mutation costs thus for their quantitative prediction. On the other hand, the degree itself of structural conservation is rather low due to the heterogeneity of the shear-design MSA: the conformation, precise location and size of the shear path, hence the role of each link, can vary from architecture to architecture, leading to low structural conservation (with peaks only around the active and allosteric site). Conservation is found much higher *within* one set of dynamically related solutions (as for Fig. 2A), corresponding to one realization of the shear design among the many included in the MSA.

## Acknowledgment

We acknowledge interesting and stimulating discussions with Eric Aurell, John Barton, Johannes Berg, Simona Cocco, Paolo de Los Rios, Solange Flatt, Joachim Krug, Michael Lassig, Duccio Malinverni, Simone Pompei, Remi Monasson, Martin Weigt, Le Yan, Stefano Zamuner. We are particularly grateful to John Barton, Le Yan, Duccio Malinverni and Stefano Zamuner for help with the codes. M. W. thanks the Swiss National Science Foundation for support under Grant No. 200021-165509 and the Simons Foundation Grant (454953 Matthieu Wyart).

---

**Figure S1:**
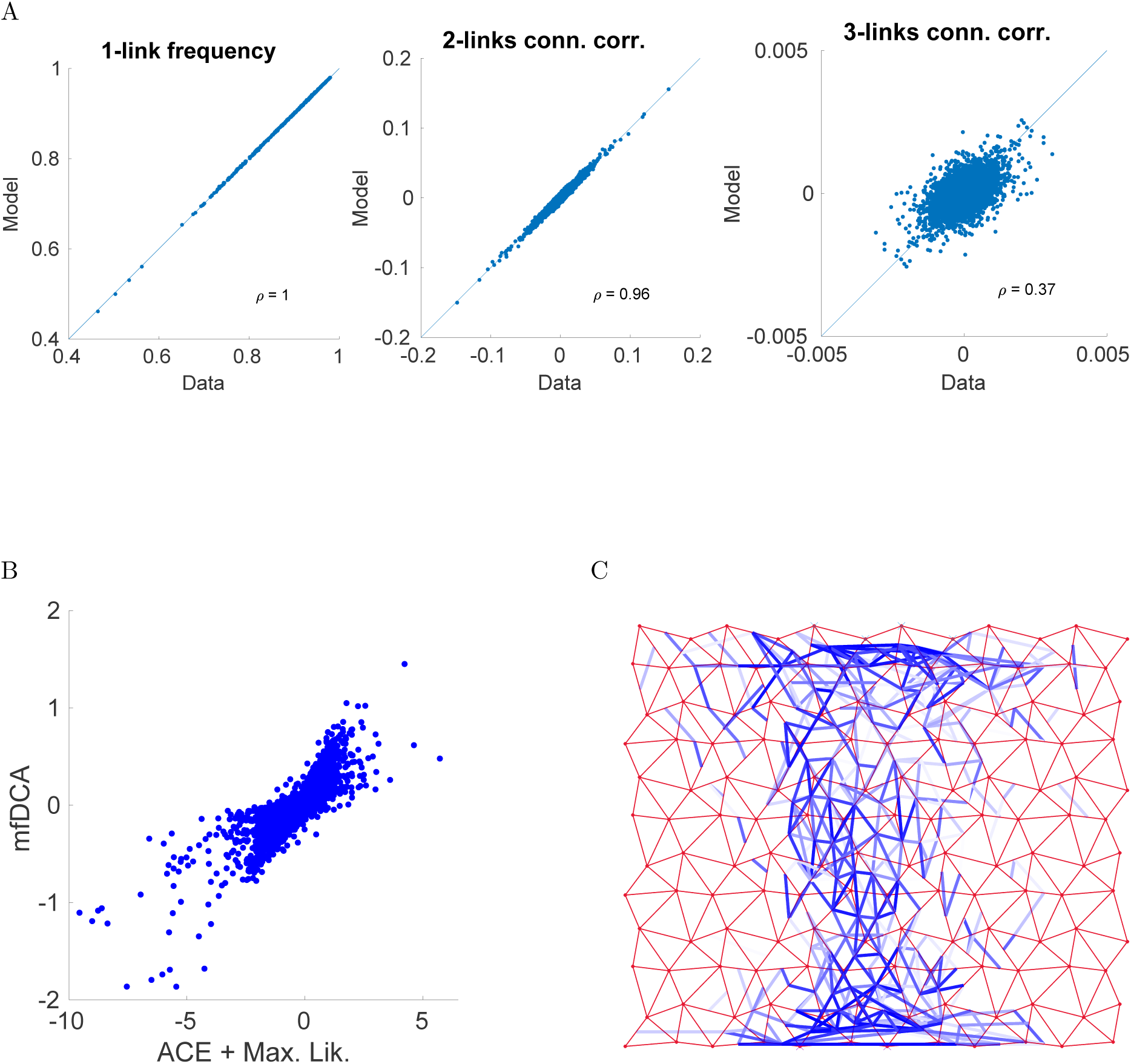
Performance of the inference procedure. A: Statistics of the model inferred by combining ACE and Maximum Likelihood. 1-link frequency and 2-links connected correlations are very accurately reproduced, as they should by construction (the relative errors, defined as in [1], are respectively *∈_m_* = 2.45 *×* 10^−1^ and *∈_C_* = 1.30 *×* 10^−1^). In contrast the third order connected correlations, which are not constrained in the inference, are not well captured (Pearson coefficient *ρ* = 0.37). This is a hint that the Ising model - a pairwise probabilistic model over *σ_i_* - is an approximation which becomes poor for estimating higher order moments. B: Scatter plot comparing *J_ij_* inferred via mfDCA to the direct couplings of ACE + Max. Lik.: the pseudocount in mfDCA has been set to *λ* = 0.5 in such a way as to obtain the highest correlation between the two. C: Spatial distribution of top 400 mfDCA-inferred couplings on the network. The reconstruction of the topology of relevant couplings is rather robust with respect to the choice of more approximate inference methods as mfDCA. As in Fig. 5A (inset) of the main text, they are concentrated at short range, i.e. they connect links lying close either to the active site or the allosteric site and in the central high-shear path. Long range mfDCA couplings, connecting links around respectively allosteric and active site, are weaker and appear among the top 600-1000 ones, implying an even worse performance at predicting long range epistasis than ACE + Max. Lik.

**Figure S2:**
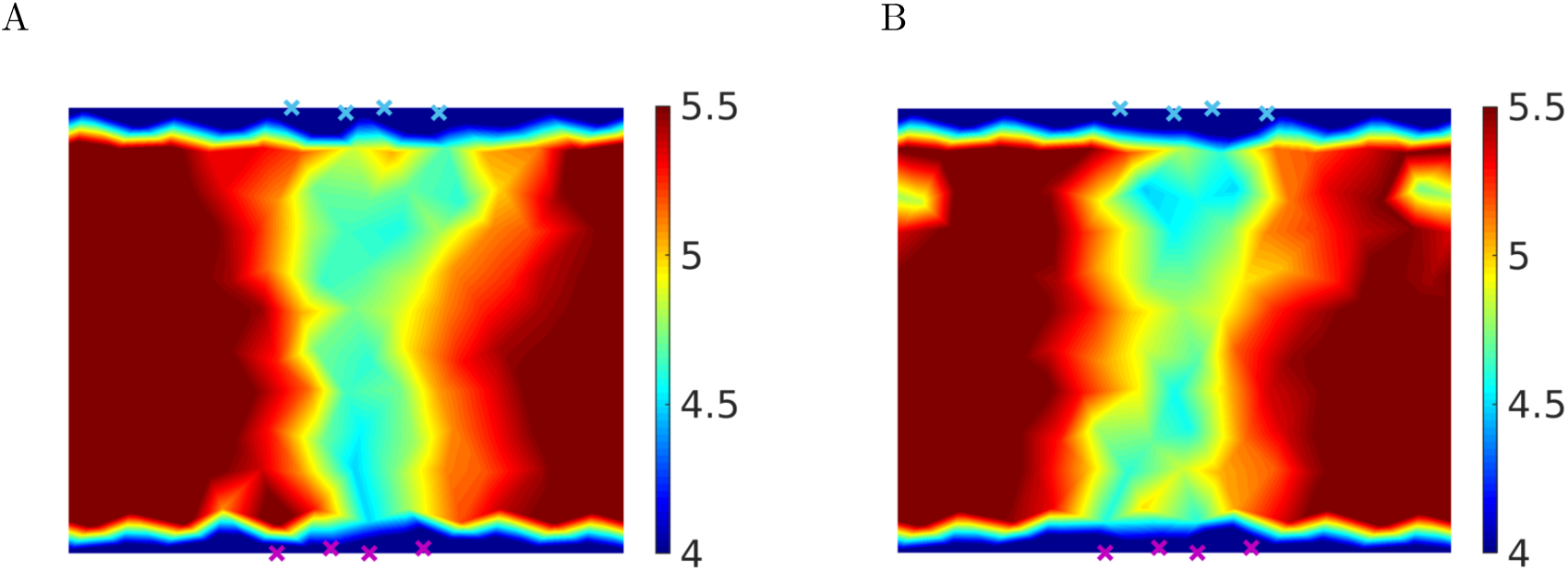
Properties of generated allosteric sequences. Coordination map of original sequences (A) and generated ones (B). They both exhibit a softer (i.e. with coordination *z <* 5) central path joining active and allosteric sites (indicated respectively by blue and purple crosses) along which the shear-like sliding takes place. This path is embedded in a more connected, “rigid” region where the coordination *z >* 5. Solutions sampled from the inferred energy landscape have a design but are not maximally fit, showing that more “structural” components, as the distribution of links, are captured but additional information would be needed to reproduce a complex mechanical function as the cooperative fitness.

## 1 Mechanical interpretation of mutation costs and epistasis

Let us denote by ε the set of nodes where ligand binding takes place, e.g. for ligand binding at the allosteric site ε = (*𝒜l*) with size dim(ε) = *n*_0_. Such event imposes a displacement ***R**^ε^* on the nodes *E* which imparts locally a force ***F** ^ε^* and induces a response ***R**^ε→r^* on all the other nodes *r*. Clearly dim(*ε*) + dim(*r*) = *L^d^* where *L^d^* is the total number of nodes for a network of size *L* in *d* dimensions; for the example of binding to the allosteric site *r* = (*𝒜c, b*), where *b* stands for the “bulk” of nodes not belonging neither to the allosteric nor to the active site. (In this paper we consider networks as in Fig. 1A of the main text, with *d* = 2, *L* = 12 and *n*_0_ = 4 for both active and allosteric site). The relation between force and overall response field is written in terms of the dynamical matrix 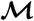

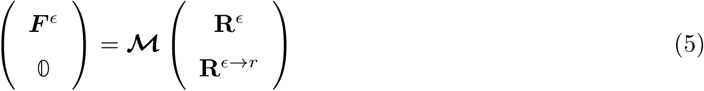

hence 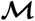 is endowed with a block structure as follows

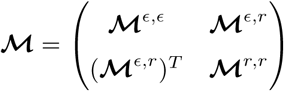

Forces as well as responses can be calculated solely from the imposed displacement by introducing a matrix 𝓠

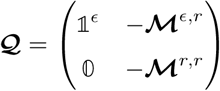

such that

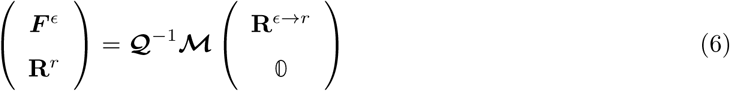

Binding at ε costs an elastic energy *E^ε^*

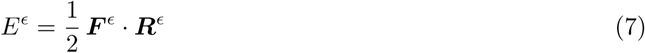

and the cooperative fitness is specified by a combination of such elastic energies

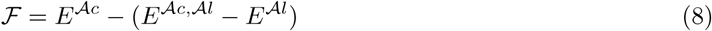

where *E^𝒜c^*, *E^𝒜c,Al^* and *E^𝒜l^* are given by Eq. 7 with *ε* = (*𝒜c*), *ε* = (*𝒜c, 𝒜l*) and *ε* = (*𝒜l*) respectively. Maximal cooperativity corresponds to making binding of a substrate at the active site energetically favored when already a ligand is bound to the allosteric site, as this reduces its binding energy from *E^𝒜c^* to (*E^𝒜c,𝒜l^ − E^𝒜l^*). One can express the energy of joint binding at the allosteric and active site 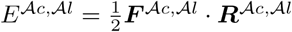 as

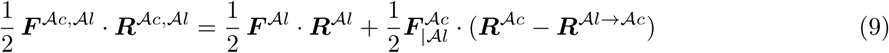

i.e. after binding at the allosteric site with an energy cost 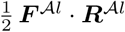, the elastic energy of binding at the active site is determined by (i) the force there when a ligand is already bound at the allosteric site (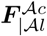 with subindex | *𝒜l*); (ii) the displacement imposed at the active site ***R**^𝒜c^* to which we subtract the response already caused by ligand binding at the allosteric site ***R**^𝒜l→𝒜c^*. Eq. 9 allows us to rewrite Eq. 8 as

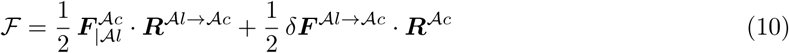

where one has 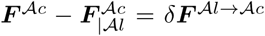. If we express *δ**F** ^𝒜l→𝒜c^* and ***R**^𝒜l→𝒜c^* in terms of the imposed displacements by using Eq. 6 and if we assume weak elastic coupling between allosteric and active site, we find that each term in Eq. 10 scales in the same way as

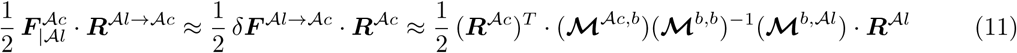

Hence, by using that 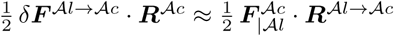, we obtain from Eq. 10

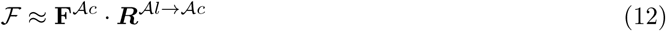

since 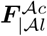 can be approximated by ***F** ^𝒜c^* in the weak coupling limit.

If we denote by 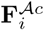 and 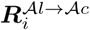 forces and displacements after a mutation at link *i*, the cost of one mutation can be expressed approximatively (see Fig. S3B) as ∆*F_i_ ≈* ∆(***F**^𝒜c^ ⋅ **R**^𝒜l→𝒜c^*)_*i*_, where 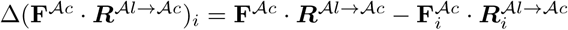. This can be further rewritten as

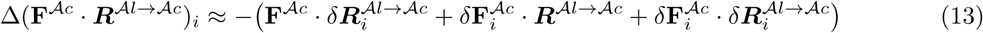

having defined changes in force as 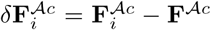 in analogy to changes in displacement 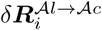 introduced in the main text. We find numerically that the cost of single mutations, when it is not too small, is dominated by the changes in displacement at the active site

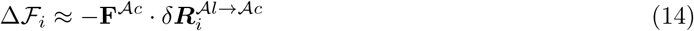

as shown in Fig. S3C. As a consequence, epistasis between mutations at *i* and *j* with significant magnitude can be written 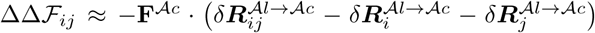, as presented in the main text. Displacement vectors and their changes upon high-cost mutations at the active site are schematically depicted in Fig. S3A.

**Figure S3:**
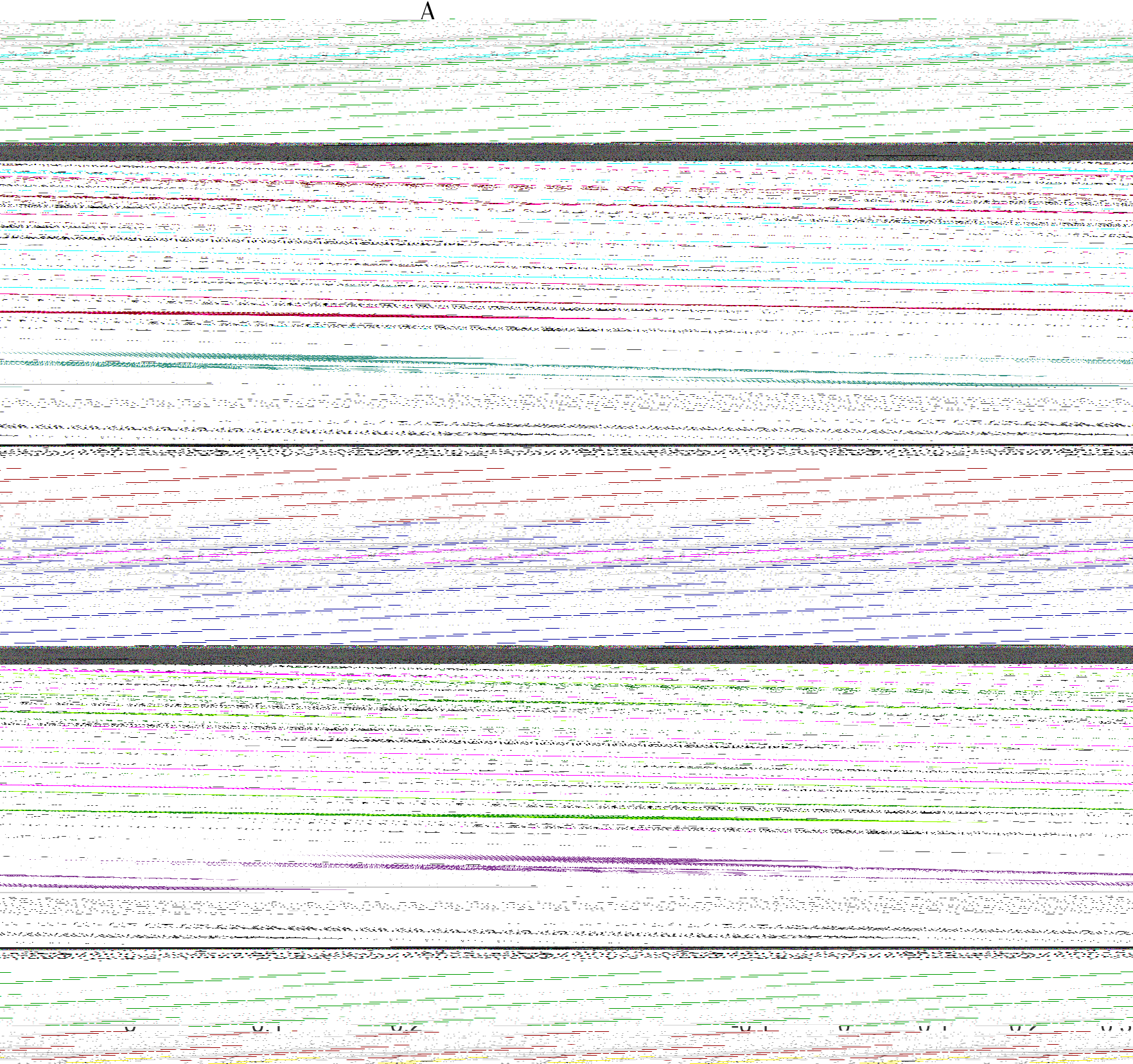
Mechanics of mutations. A: The geometry of mutation costs is illustrated in the zoom on the active site region (note that for simplicity of visualization we consider only one of the *n*_0_ = 4 nodes). Thick, dark red lines highlight links whose disruption would be lethal for the allosteric fitness. These few links, crucial to the long-distance propagation of the allosteric response, are located around active and allosteric site and exhibit maximal epistasis along with maximal single mutation costs (i.e. they populate the saturation region of Fig. 2A in the main text). After a lethal mutation consisting in removing a spring at link *i*, the displacement at the active site 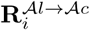 is significantly reduced with respect to the original optimal displacement ***R**^𝒜l→𝒜c^* and their difference is given by 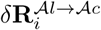 (dashed arrow). When a second lethal mutation at *j* occurs, we denote by *θ* the angle between 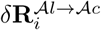 and 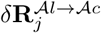; for lethal mutations cos(*θ*) *≈* 1 (see Fig. 2B in the main text), i.e. they all tend to have a homogeneous direction of action which is precisely the one opposite to the displacement at the active site. B: Numerical test of the approximation ∆*𝓕_i_ ≈* ∆(***F**^𝒜c^ ⋅ **R**^𝒜l→𝒜c^*)_*i*_ and of 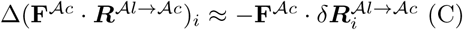. The latter is valid only for medium-high mutation costs.

## 2 Prediction of epistasis

The scaling of epistasis (Eq. 2 in the main text) suggests a measure simply based on the inferred single mutation costs, i.e. *|* ∆∆*𝓕_ij_| ∝* min(∆*𝓔_i_,* ∆*𝓔_j_*). We have verified that this improves extremely the prediction of long-range epistasis in our model for allostery, both for single configurations and for the average epistatic pattern, as shown in respectively in Fig. S4B and C. The measure of epistasis via top *| J_ij_|* requires the inferred model to be performant at capturing local information via local parameters; on the other hand, the estimation of single mutation costs incorporates all the local parameters inferred from the statistics. These results support the view that more functional information (related to non-local modes) is embedded in weaker couplings which would be excluded by applying the contact-prediction criterion of looking at the largest ones (usually as many as the system size): for example recently [2] has found that the prediction of functional cooperativity between distant sites could be improved by considering several “non-contacting” DCA couplings.

### 2.1 Simple model illustrating the failure of DCA

To explain the discrepancy between short-range and long-range DCA-predictions of epistasis, we resort to the simple model of Fig. 6 (main text). We assign to all the 49 functional configurations the same fitness *𝓕*, all the other 2^8^ *−* 49 configurations would not belong to the sample of optimal configurations and are taken with zero fitness, thus ∆*𝓕*= 0 if a mutation (single or double) results in a configuration still belonging to the optimal sample and ∆*𝓕*= *𝓕*otherwise. We can estimate average mutation costs by counting how frequently mutations would lead to a configuration outside of the optimal sample, yielding

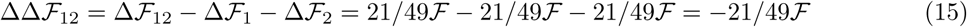

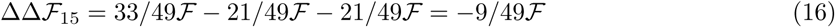

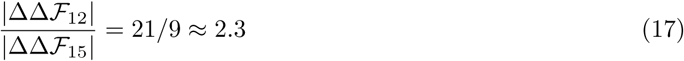

Next, by a simple likelihood maximization we infer the set of *J_ij_* and *h_i_* compatible with *(σ_i_)* and *(σ_i_σ_j_)*, single-site and pairwise frequencies of the optimal sample. We estimate *J*12 = 1.18 and *J*15 = 0.40, thus the prediction by DCA

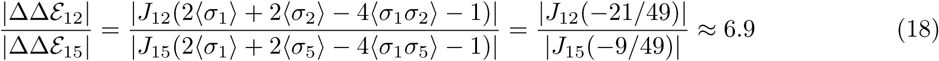

i.e. the DCA prediction is significantly biased towards short-range epistasis. Due to symmetry of our model, epistasis and the DCA-prediction for any combination of units in the two subparts is the same as for units 1 and 5; similarly, the result for 2 units within the same group is given by the values for units 1 and 2. For the remaining combinations of units, i.e. the ones belonging the same subpart but to different groups (e.g. *i* = 1 and *j* = 3) we obtain that epistasis is weaker compared to units within the same group

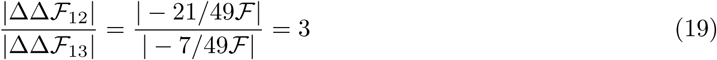

**Figure S4:**
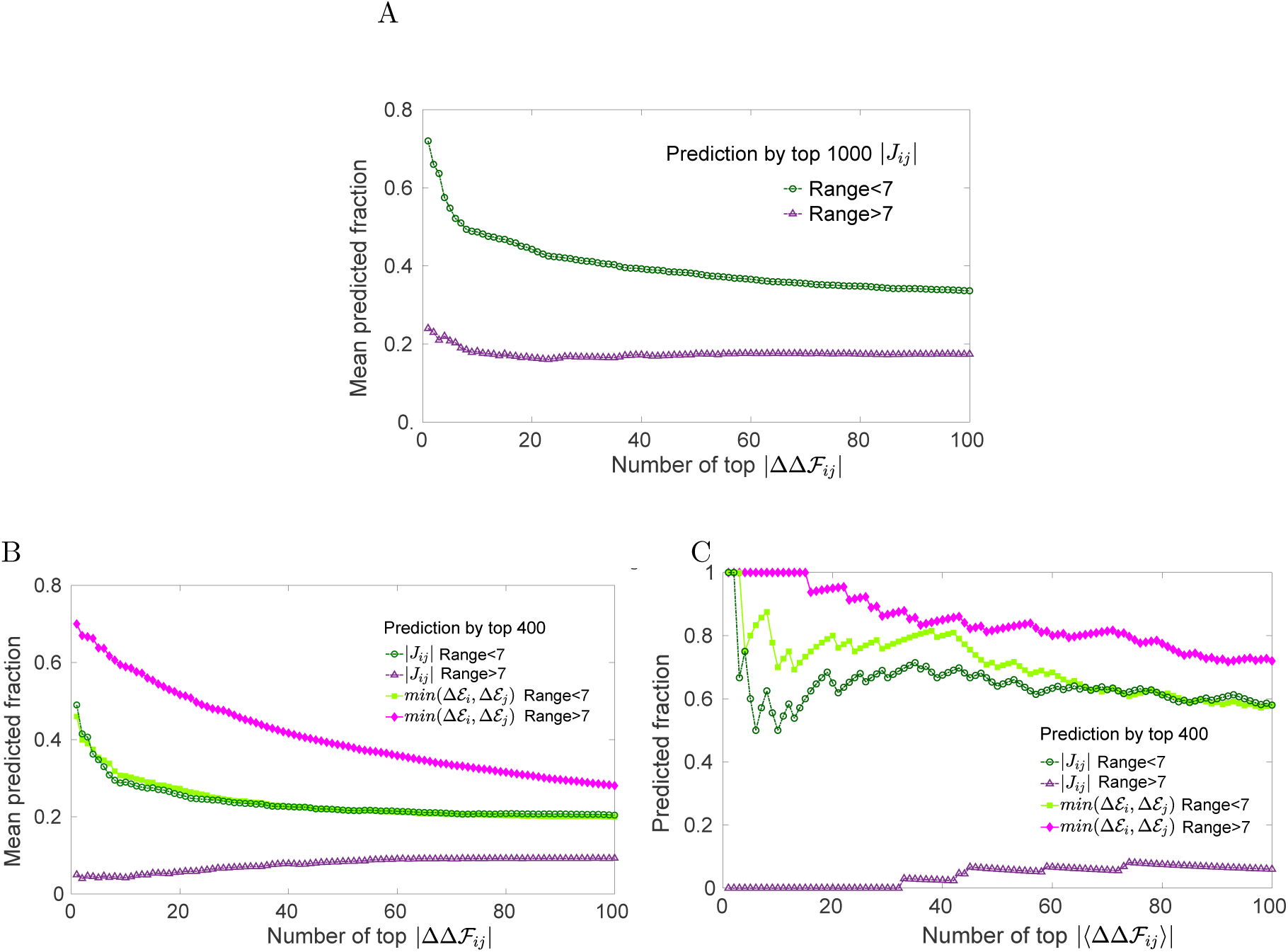
Prediction of epistasis by the DCA-inferred model. A: Same plot as in Fig. 5B (main text) where we show the fraction of top rank epistasis *|* ∆∆*𝓕_ij_|* predicted by top 1000 *| J_ij_|*, averaged over 100 configurations. In comparison to Fig. 5B, here we consider a higher number of the largest in magnitude couplings to predict epistasis: the mean predicted fraction increases both for short range and long range epistasis, yet a clear difference between their values remains. B: Same plot as Fig. 5B (main text) where we added curves for the prediction by min(∆*𝓔_i_,* ∆*𝓔_j_*) - the minimum between average single mutation costs at *i* and *j* - as implied by scaling 2 in the main text. As in Fig. 5B, we rank separately long-range (> 7) and short-range (< 7) pairs of links *i* and *j* in terms of *|* ∆∆*𝓕_ij_|* and we plot the fraction of these pairs - averaged over 100 configurations randomly chosen - falling either into the top 400 *| J_ij_|* (empty symbols) or into the top 400 values of min(∆*𝓔_i_,* ∆*𝓔_j_*) (filled symbols). This second measure improves only slightly the estimation of strong short-range epistasis but it does so dramatically for long-range one. C: Same plot as B where we show the fraction of the average epistasis (∆∆*𝓕_ij_)* (estimated from 1.5 *×* 10^3^ randomly chosen configurations of the MSA) that one would predict either via *| J_ij_|* or min(∆*𝓔_i_,* ∆*𝓔_j_*). The prediction at short distance is rather accurate, with the predicted fraction reaching 1 for the maximally epistatic pairs; at long distance, signal on long-range epistasis captured by *| J_ij_|* is almost absent while the prediction by min(∆*𝓔_i_,* ∆*𝓔_j_*) stands out for its precision.

Since each subpart can be of different type (OR gate), units from different groups (i.e. types) are less tightly constrained by function. The DCA-prediction does not underestimate epistasis as for units of different subparts (i.e. at long distance) with

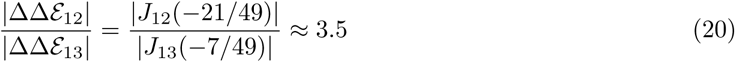

where *J*_13_ = *−*1.01. From Eq. 17, Eq. 18, Eq. 19 and Eq. 20 it is straightforward to calculate *|* ∆∆*𝓔*_13_*| /|* ∆∆*𝓔*_12_*| × |* ∆∆*𝓕*_12_*| /|* ∆∆*𝓕*_13_*| ≈* 0.86 and *|* ∆∆*𝓔*_15_*| /|* ∆∆*𝓔*_12_*| × |* ∆∆*𝓕*_12_*| /|* ∆∆*𝓕*_15_*| ≈* 0.33

#### 2.1.1 Feedforward neural network

To understand which machine learning tools could improve the prediction of epistasis in the simple model, we have built a feedforward neural network performing least squares regression of sequence data based on their fitness (see Fig. S5). For data in the training set, we provide the network with both the input sequence and the target answer, i.e. a label 1 (standing for fitness 𝓕) or 0. We vary the size of the training set from 50% to 80% of the 2^8^ = 256 total sequences and we keep 20% of the sample for validation of the accuracy of prediction. We learn the weights, i.e. the connections between layers, which minimize the mean squared error between the output of the network and the target answers by stochastic gradient descent from a random orthogonal initialization; only relatively few trainings (about 1 in 10) find a high quality solution. We obtain that the mean squared error between true and estimated fitness, averaged over 100 of such high-quality trainings, ranges between *∼* 2 *×* 10^−6^ for a training set with 50% of the sample to *∼* 2 *×* 10^−7^ with 80%. Therefore, when the network is presented with an optimal sequence mutated at some position, the network can predict the value of its fitness with extreme accuracy in such a way as to predict ∆*𝓕 ∼* 0 when it still belongs to the optimal sample or ∆*𝓕 ∼* 1 if it does not. This ensures that also epistasis would be accurately predicted at any range.

---

**Figure S5:**
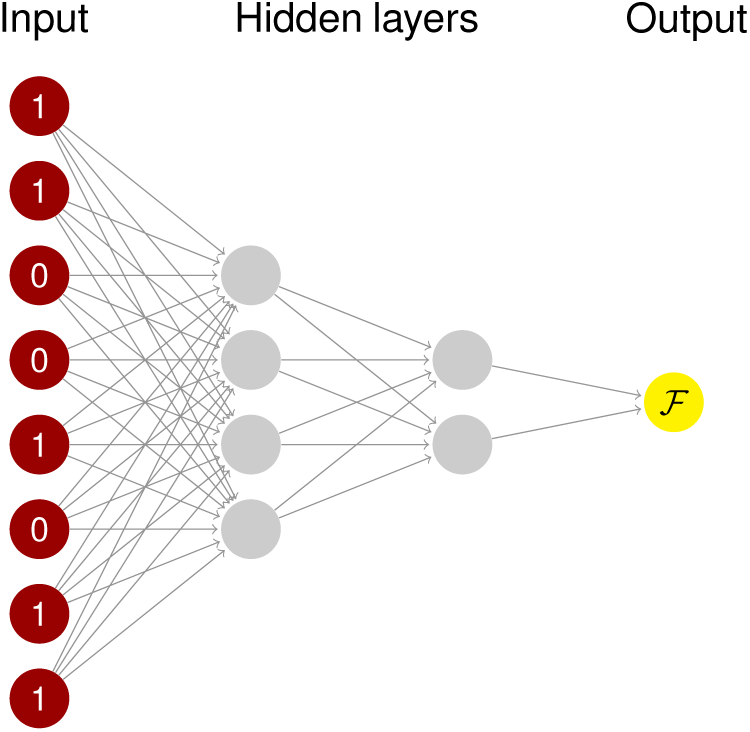
Graphical representation of the feedforward neural network for regression in the simple model. The size of the input layer is 8, as the size of the system. We add two hidden layers of 4 and 2 units and the final one-unit output is 1 if the input sequence has fitness *𝓕* and 0 otherwise. The activation function from one layer to the successive one is a sigmoid and the weights are dense (all units in one layer are connected to all units of the successive one).

